# Rapid turnover and evolution of sex-determining regions among *Sebastes* rockfishes

**DOI:** 10.1101/2023.01.19.524770

**Authors:** Nathan T.B. Sykes, Sree Rohit Raj Kolora, Peter H. Sudmant, Gregory L. Owens

## Abstract

Nature has evolved a wealth of sex determination (SD) mechanisms, driven by both genetic and environmental factors. Recent studies of SD in fishes have shown that not all taxa fit the classic paradigm of sex chromosome evolution and diverse SD methods can be found even among closely related species. Here, we apply a suite of genomic approaches to investigate sex-biased genomic variation in eight species of *Sebastes* rockfish found in the northeast Pacific Ocean. Using recently assembled chromosome-level rockfish genomes, we leverage published sequence data to identify disparate sex chromosomes and sex-biased loci in five species. We identify two putative male sex chromosomes in *S. diaconus*, a single putative sex chromosome in the sibling species *S. carnatus* and *S. chrysomelas*, and an unplaced sex determining contig in the sibling species *S. miniatus* and *S. crocotulus*. Our study provides evidence for disparate means of sex determination within a recently diverged set of species, and sheds light on the diverse origins of sex determination mechanisms present in the animal kingdom.

## Introduction

Sexual reproduction is common to all higher order animals, and nature has evolved a great diversity of mechanisms to produce distinct sexes, both within and among animal taxa. The mechanisms underlying sex determination (SD) may be genetic (GSD), environmental (ESD), or some combination of both (Bachtrog et al., 2014). Even among organisms with GSD, there is substantial diversity owing to the many independent origins of sex chromosomes across the tree of life (Renner & Ricklefs, 1995; Mank et al., 2006; Pokorná & Kratochvíl, 2009). Most familiar among GSD mechanisms is heterogametic sex determination (HSD), in which the presence or absence of a particular sex chromosome drives gonadal differentiation. While eutherian mammalian males are heterogametic (XY), in birds and moths (among many others) females are the heterogametic sex (ZW), whereas males are homogametic. These systems have become nearly fixed in their clades, giving the impression of HSD as an ultimate and inevitable outcome of sex chromosome evolution. Conversely, the fusion of a depleted mammalian Y-chromosome to the X in creeping voles highlights the ephemeral nature of SD mechanisms (Couger et al., 2021). Indeed, many taxa possess sex-determining regions or chromosomes which are not fixed, and which turn over frequently on an evolutionary time scale. This lability allows us to understand the origin of SD systems by examining sex determining regions at different stages of differentiation.

The classical philosophy surrounding the evolution of HSD suggests a logical series of steps (Muller, 1964; Ohno, 1967; Kratochvil et al., 2021). First among these is the acquisition of a master sex determining (MSD) gene, an allele which may itself play a key role is testis differentiation or modulate expression at other loci. However, to evolve from a single mutational difference between sexes to highly differentiated sex chromosomes, recombination between haplotypes containing differing sex determining alleles needs to be suppressed. Without this second step, any linkage between additional mutations and the MSD, including sexually antagonistic loci, will inevitably decay. Without recombination suppression, genetic differentiation between sexes would remain limited to one or more mutations that directly control sex, rather than a whole sex-determining region.

There are several models for how recombination suppression can evolve and what consequences might ensue (Wright et al., 2016). For instance, the sexual antagonism model suggests selection against recombination around the MSD and linked genes with sex-specific effects leads to sex-specific supergenes. These gene complexes are passed to offspring as a unit, who then simultaneously inherit a suite of tightly linked, sex-specific adaptations. While the sexual antagonism model is broadly supported by theory (Rice, 1987; van Doorn & Kirkpatrick, 2007) and evidence (Rice, 1992; Gibson et al., 2002; Zhou & Bachtrog, 2012), Ponnikas et al. (2018) note that recombination suppression may be overly attributed to such conflict. Additional mechanisms and rationale for recombination suppression in young sex chromosomes have been proposed. These include: (1) selfish elements within X chromosomes (Jaenike, 2001), which distort sex ratios by supressing or eliminating Y-linked alleles (2) heterozygote advantage (Charlesworth & Wall, 1999), whereby individual fitness is improved by masking deleterious recessive mutations in non-recombining regions and (3) random accumulation of mutations or chromosomal structural variants (genetic drift; Charlesworth et al., 1987). Another plausible explanation for reduced recombination involves mutations to transcription factor binding sequences, outside the protein-coding region, which can rapidly result in highly specific heterochiasmy (i.e., sex-specific recombination) within populations (Kong et al., 2010). Whatever the mechanism of recombination suppression, the differentiating, heterogametic sex chromosome exhibits weakened selection against deleterious alleles due to reduced recombination.

The accumulation of mildly deleterious alleles can deactivate genes on the Y chromosome, leading to reduced positive selection in maintaining those regions and an overall size reduction through the Y-degeneration process (Bachtrog, 2013). This one-way trajectory can result in a relatively stable sex chromosome over short evolutionary time but leaves a degenerated chromosome vulnerable to loss from the genome over longer timescales, as was the case with creeping voles. One model to explain this loss is Muller’s ratchet, summarized by Charlesworth & Charlesworth (2000) as a stepwise, stochastic loss of the Y chromosomes carrying the least deleterious load, in which each loss results in the fixation of one or more deleterious alleles in the population (Muller, 1932). As a result, the Y chromosomes in circulation within the population become increasingly impaired in their function. This mechanism is especially plausible in small populations but may also act over relatively short evolutionary time even when populations are large (Gessler, 1995).

While sex determining regions are highly variable, MSD genes are often convergent and frequently involved in testis differentiation (Marshall Graves & Peichel, 2010). Indeed, a ‘limited options’ hypothesis has been proposed to explain the convergence of SD genes among vertebrates, owing to the small number of players in this role. Among these are the androgen receptor (*ar*), Anti-Mullerian hormone (*amh*), Doublesex and mab-3 related transcription factor (dmrt1), and *sox*-*3*/*SRY* genes (Kratochvil et al., 2021; Marshall Graves & Peichel, 2010). Their critical role in gonadal development makes these genes susceptible to cooption as MSD, as rare gain-of-function mutations can easily tip the scales by promoting differentiation to one sex or disabling the other.

The classical model fits taxa with relatively ancient and conserved HSD but a growing body of evidence in other taxa, without fixed SD, points to a higher flexibility in the evolution of sex chromosomes (Kratochvil et al., 2021; Li & Gui, 2018). Among vertebrates, teleost fishes exhibit the most variability in sex-determination mechanisms (Bachtrog, 2014). Among these are ESD, as in many flatfishes (Luckenbach et al., 2009), socially or environmentally driven sequential hermaphroditism in gobies (Sunobe et al., 2017), variable MSD in several clades including salmonids, halibut, and tuna (McKinney et al., 2020; Chiba et al., 2021; Edvardsen et al., 2022).

The strongest evidence for lability in SD regions can be found in taxa with newly or only slightly differentiated sex chromosomes. One such example was recently discovered in cichlid fishes (El Taher et al., 2021), whose SD locus is highly labile on short evolutionary timescales. Cichlids comprise an estimated 3000 species, over 200 of which underwent analysis for SD mechanisms in the study by El Taher et al. (2021). The study found evidence for frequent turnovers in poorly differentiated sex chromosomes, sex chromosome fusions and other large-scale rearrangements, as well as some convergent and well-conserved sex-linkages in some clades. The highly variable origin, breadth, and position of these sex-linked regions within cichlids demonstrates the ephemeral nature of SD in taxa without a fixed mechanism.

*Sebastes*, a highly speciose genus of rockfish, underwent rapid speciation in the Pacific Ocean (Mangel et al., 2007). Among northwest Pacific rockfish, three species have been found to possess a duplicated *amh*, acting as an MSD gene (Song et al., 2021). The redundancy conferred by a gene duplication is a prime candidate for neofunctionalization (Edgecombe et al., 2021). Several independent duplications of *amh* among teleost fish have indeed acquired sex-determining functions (Hattori et al., 2013), examples include tilapia (Liu et al., 2022), stickleback (Jeffries et al., 2022), and ayu (Nakamoto et al., 2021). The study by Song et al. (2021) notably failed to attribute the same MSD mechanism to a handful of northeast Pacific rockfish, indicative of variable SD systems within a single, recently speciated genus.

Recent genome sequencing of over 80 species of Pacific rockfish (Kolora et al., 2021) permits chromosome mapping and analysis of previously published sequence data. Here, we leverage previously published Restriction site Associated DNA sequencing (RADseq) data for eight species of Pacific rockfish to identify putative regions responsible for SD. We make use of a novel high throughput approach, as well as traditional variant calling methods, to identify genomic variation associated with sex. Additionally, we assemble a phylogeny of whole gene sequences for *amh* in Pacific rockfish including 78 *Sebastes* spp. and close relatives, to place the duplication in evolutionary context and identify species for further investigation of *amh* as the MSD gene.

## Materials & Methods

### Data acquisition

Previously published RADseq data was publicly available from NCBI for eight species of *Sebastes* (Tbl. 1). These included the gopher (*S. carnatus*) and black-and-yellow (*S. chrysomelas*) rockfishes (Fowler & Buonaccorsi, 2016), sunset (*S. crocotulus*) and vermillion (*S. miniatus*) rockfishes (Longo et al., 2021), boccacio (*S. paucispinis*), canary (*S. pinniger*) and yelloweye (*S. ruberrimus*) rockfishes (Andrews et al., 2018) and deacon (*S. diaconus*) rockfish (Vaux et al., 2019).

**Table 1.**
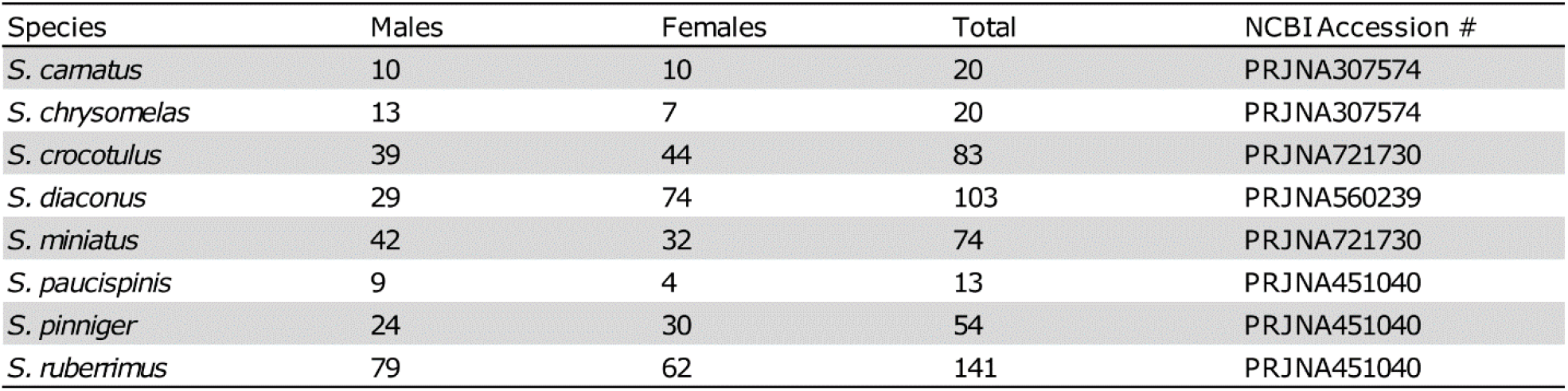
Summary of sample species. Our study comprises eight total species from four previously published datasets. Sample sizes and sex ratios varied between species.

As our analyses relied on sexed sequence data, unsexed samples were omitted. Additionally, the admixture analysis of sunset and vermillion rockfish, from which our data was sourced, found evidence of several putative hybrid individuals in their dataset, which were also omitted from our analysis.

### GWAS for sex bias

To identify sex-biased markers, we first used RADSex (Feron et al., 2021; RADSex version 1.2.0). RADSex compares identical RADseq reads to assess presence or absence of markers in each individual and calculates the number and distribution of markers based on read depth.

Unaligned sequences in FASTQ format were fed into RADSex alongside a table containing sample accession numbers and sex, with Bonferroni statistical correction disabled. Yates’ correction for continuity was left in place for all analyses, as some species had low sample numbers (n < 30). After identification, markers were mapped to species-specific reference genomes, when available (Kolora et al., 2021). In the cases of *S. crocotulus* and *S. chrysomelas*, for whom genomes were unavailable, markers were mapped to those of their sibling species, *S. miniatus* (diverged <5Mya) and *S. carnatus* (diverged <2.5Mya), respectively. All reference genomes used were generated using Illumina sequence data and scaffolded against the chromosome-level *S. aleutianus* genome using RagTag (Alonge et al., 2021). This common scaffolding means that positions are roughly syntenic between the genomes and allows for easy comparison.

For use in a traditional variant calling pipeline, the sequences were mapped to the above reference genomes with the Burrows-Wheeler aligner (Vasimuddin et al., 2019; bwa-mem2 version 2.2.1), read groups were appended with Picard (Broad Institute, 2019; Picard Toolkit version 2.26.3), converted to BAM and sorted with samtools (Danecek et al., 2021; samtools version 1.13).

We then used *freebayes* (Garrison & Marth, 2012; freebayes version 1.3.5) to call variants individually, producing a separate VCF (variant call file) for each scaffold, parallelized with *GNU parallel* (Tange, 2018). When calling *S. diaconus* variants, spikes in computational resource allocation required further separation of scaffolds for chromosomes 13, 19 and 20 into 5Mb subsections. All intermediate VCFs were then sequentially combined into a single file for each species.

Custom Perl scripts were used to compute variation between the sexes using three metrics: (1) presence/absence of individual loci (missing data), (2) allele frequency and (3) heterozygosity at each locus. Our scripts included Pearson’s chi-square test of independence with Yates’ correction for continuity, for a likewise comparison with RADSex outputs. All scripts used in analysis are included in a git repository (github.com/ntbsykes/rockfish_sex).

To visualize the results, we calculated the proportion of markers with a p-value < 0.005 for all test statistics across a sliding 250bp window (Fig. 1). Given our relatively small sample size and high marker number, it is unlikely for any individual marker, in some species, to reach statistical significance using traditional genome-wide association (GWA) thresholds. With the sliding window, we instead identified regions with unusually high concentrations of near-significant markers, which is indicative of sex differentiation.

**Figure 1.**
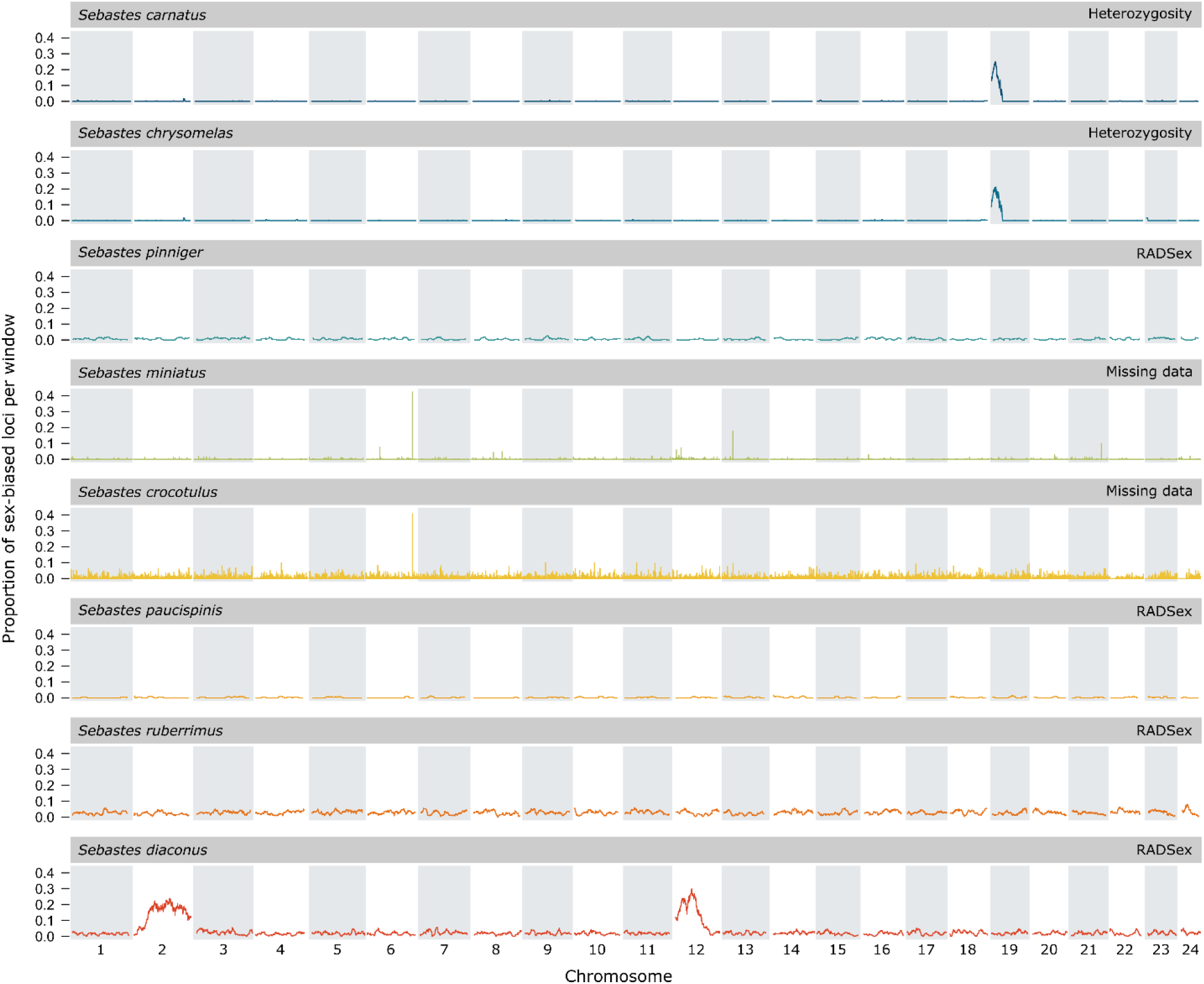
Whole genome rolling mean of significant SNPs. Markers were characterized as significant or not based on a probability threshold of association with sex (P < 0.005). Significant markers were given a value of 1, while insignificant markers were zero and a rolling mean was calculated across windows of 250 SNPs for the whole genome. For clarity, markers from the comparative metrics with the clearest signal in each species were used for visualization: *S. carnatus S. chrysomelas* – heterozygosity; *S. crocotulus S. miniatus* – missing data; *S. paucispinis, S. pinniger, S. ruberrimus S. diaconus* – RADSex.

For further investigation of two sex-determining regions in *S. diaconus*, we compiled a list of the most significant loci with a higher allele frequency in males (-log(P) ≥ 20). After isolating these sites using *vcftools* (vcftools version 1.13; Danecek et al., 2011), we used *R* (R version 4.1; R Core Team, 2021) to plot the genotype at each locus and mean number of male-biased alleles for all samples (Fig. 2).

**Figure 2.**
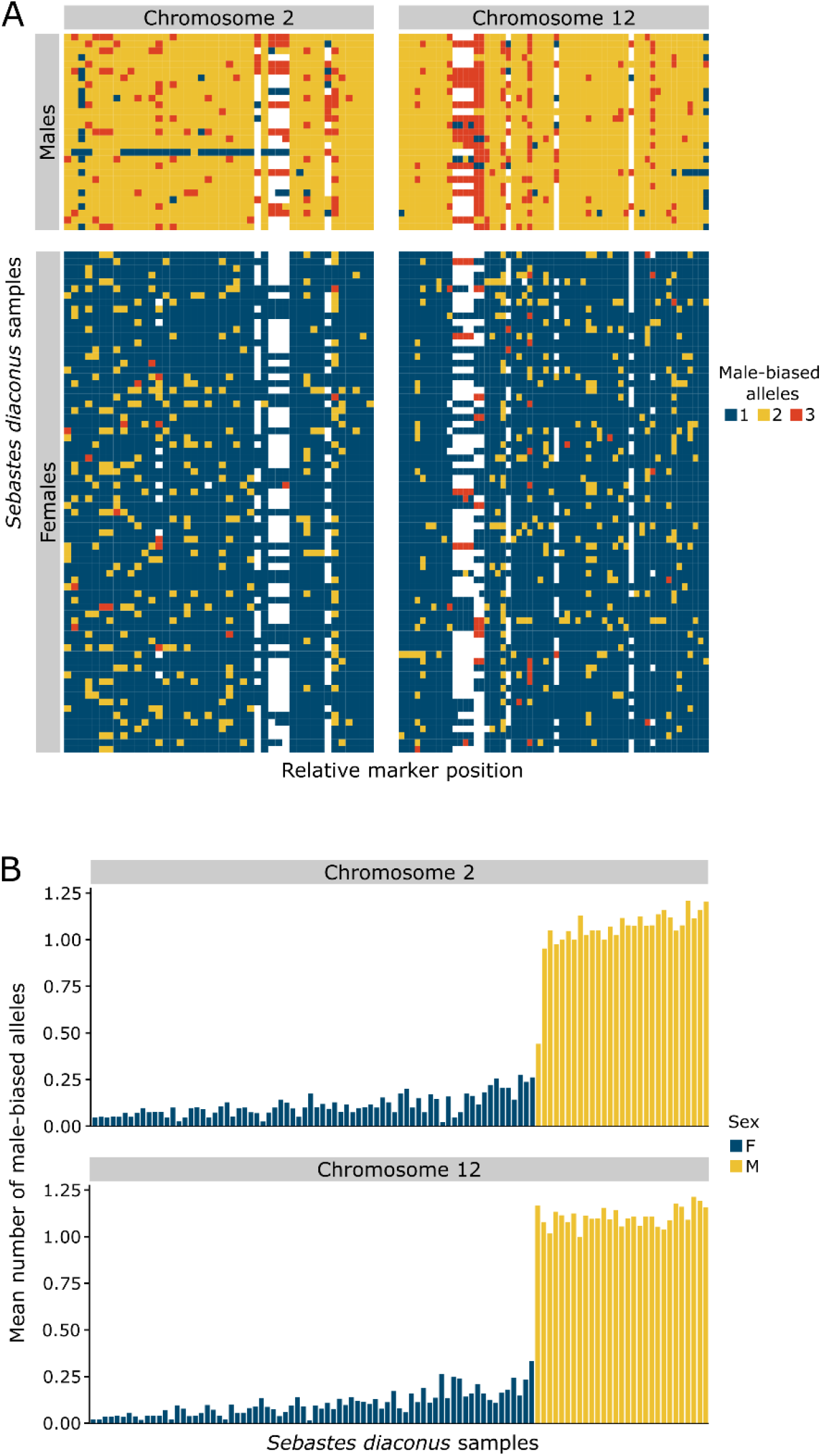
A) Comparison of per-sample male-biased alleles in *Sebastes diaconus*. Significantly male-biased alleles (-log(P) ≥ 20; each locus represented by one square and arranged by their position on the chromosome) are present on both chromosomes 2 and 12 for nearly all samples. The sex-determining locus likely lies in a region with alleles shared by all males. **B) Per-sample mean number of significantly male-biased alleles on both sex chromosomes**. Females, shown in blue, possess significantly fewer male alleles at these sites on average

### Mapping canary, sunset, and vermillion rockfish data to higher quality genomes

The initial Illumina-sequenced and RagTag-assembled genomes used for alignment provide a common order for identifying shared regions but are lower quality than *de novo* assembled long-read genomes. We leveraged two high quality reference genomes for more accurate alignment of three species’ datasets: *S. pinniger* (GCA_916701065.2,) *S. miniatus*, and *S. crocotulus* (the latter two both aligned to the *S. miniatus* genome GCA_916701275.1). We then used NCBI blastn (Camacho *et al*., 2009; blast+ version 2.12.0) to compare the nucleotide sequence of observed peaks to detected by the previous alignment. Finally, to characterize the extent of differentiation at these loci, we applied a similar approach to our isolation of sex-biased loci in *S. diaconus* (this time -log(P) ≥ 7; Fig. 3B), extracted read depth at each locus using bcftools (Danecek et al., 2021; bcftools version 1.13), and averaged the read depth by sex across the entire unplaced contig, taking the difference (Fig. 3C).

**Figure 3.**
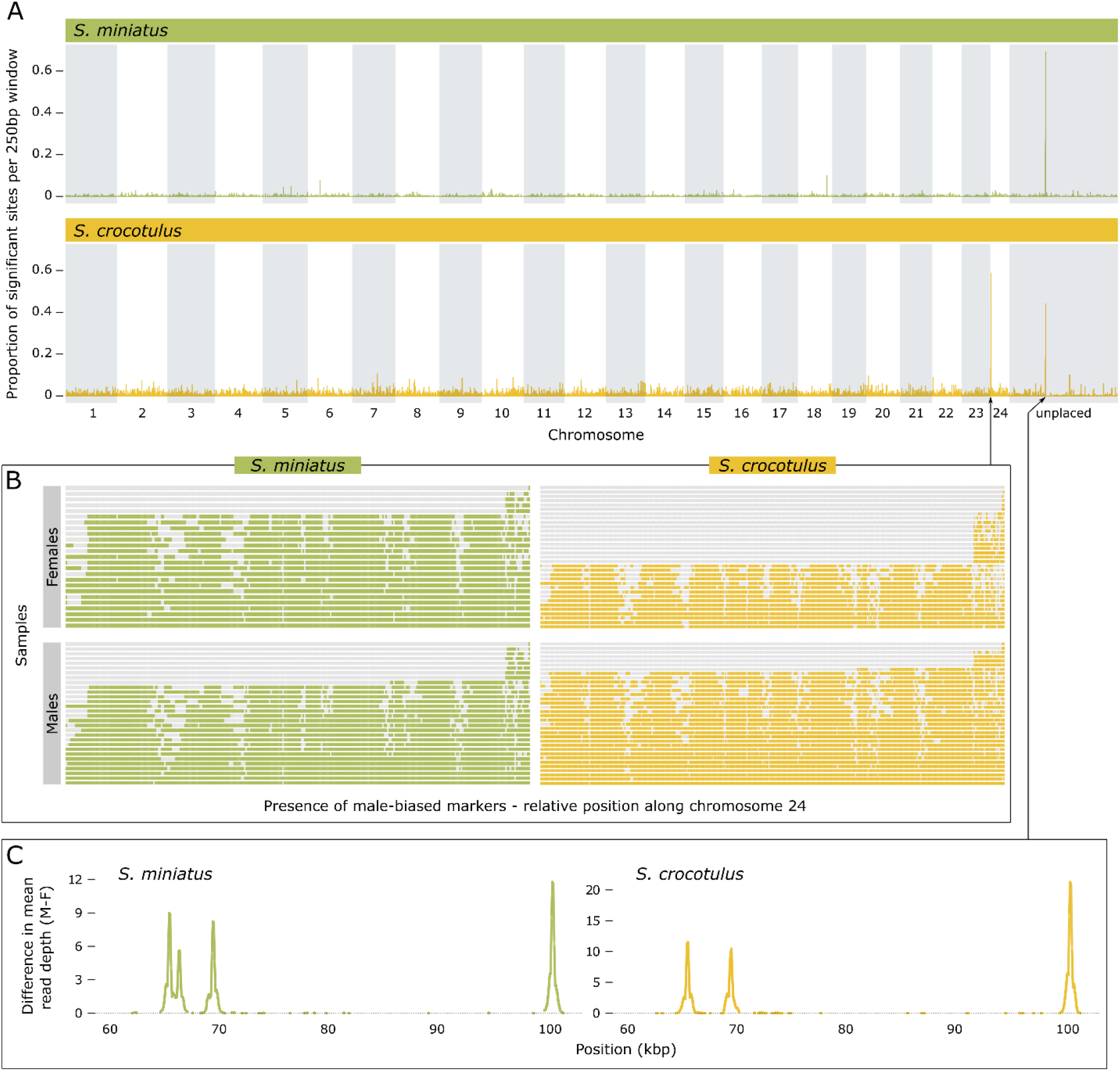
A) High quality genome alignment of *S. miniatus* and *crocotulus* reveals regions significantly associated with maleness. Both species ‘ data were mapped against the *S. miniatus* genome. One sex-biased region, shared by both species, maps to an unplaced contig. Significant sex bias was detected in *S. crocotulus* along the first megabase of chromosome 24. **B) Missing data among *S. crocotulus* females leads to significant sex bias on chromosome 24**. Distinct sample groupings may represent patterns of recombination. **C) Difference in mean read depth between the sexes along the entirety of a sex-biased, unplaced contig**. Loci are homologous to those on chromosomes 6, 13, and 17 in the Figure 1 ragtag assemblies.

### Assembly of amh phylogeny in Sebastes and their relatives

Coding sequences for *S. schlegelii amha* and *amhy* were acquired from NCBI (MW591742, MW591743) (Song et al., 2021) and queried against 88 rockfish genomes (Kolora et al., 2021) and three-spine stickleback (GAculeatus UGA version5) using blastn in command-line, yielding matches on exons. We parsed the BLAST results in R to determine gene start and end position, as well as coding strand. We then used samtools faidx to extract the whole gene sequence for all copies of *amh* in each species.

These were aligned with MUSCLE multiple sequence alignment tool (Edgar, 2003). The alignments were manually checked in BioEdit 7.2.5 (Hall, 1999). Finally, a maximum likelihood phylogeny of all gene copies was produced in iqtree 1.6.2 (Nguyen et al., 2015) using default parameters and 1000 bootstrapping replicates. The resulting gene phylogeny was visualized in Figtree (Rambaut, 2018; Fig. 4).

**Figure 4.**
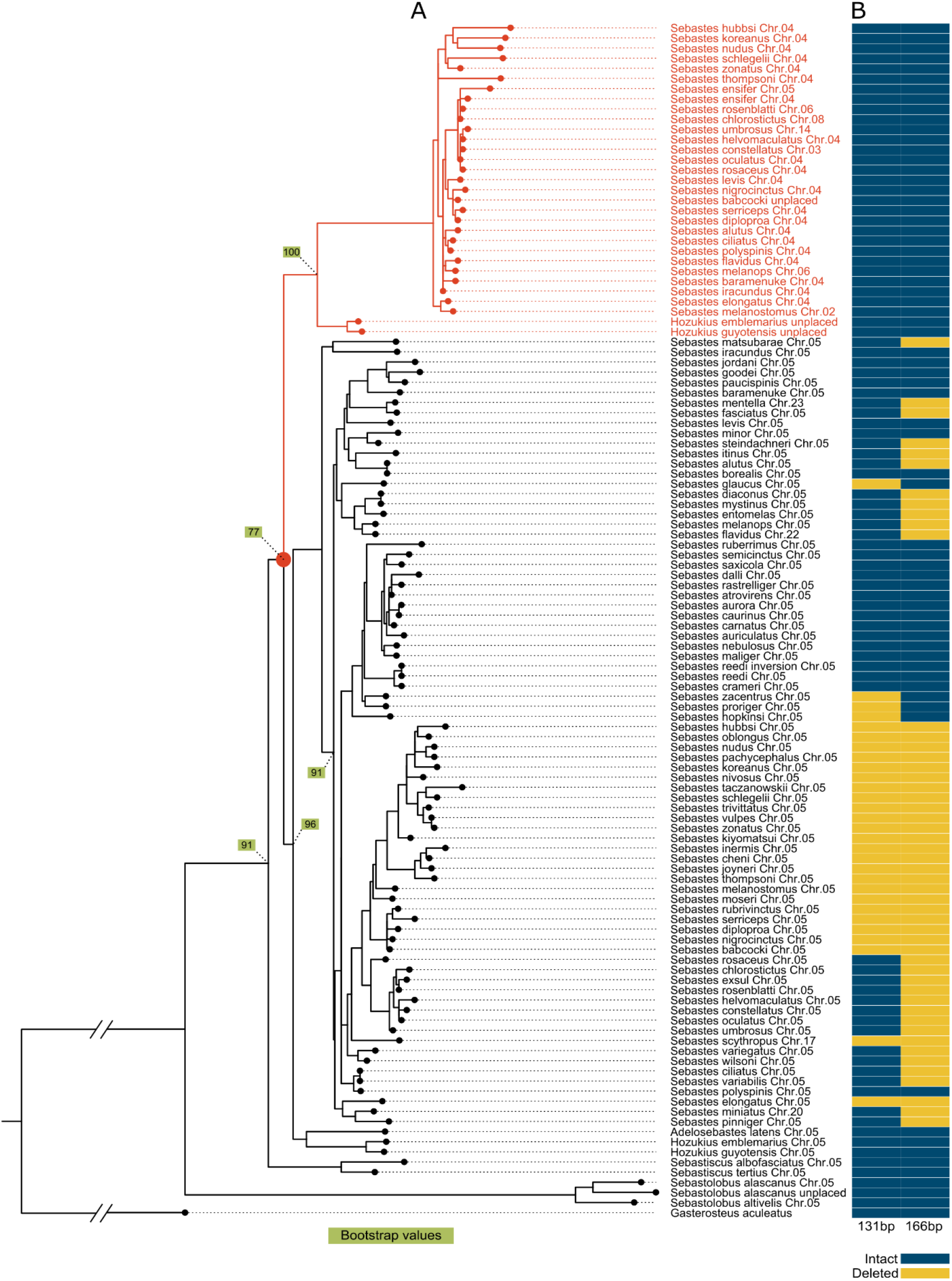
A) Maximum likelihood phylogeny for *amh* and *amhy* in *Sebastes*, with outgroups. The *amh-amhy* duplication event is shown with a red circle and *amhy* copies are highlighted in red. Bootstrap values, highlighted in green, show confidence in sorting at nodes of particular interest. **B) Table of intron 4 deletions in *amh***. Both deletions are derived in the autosomal *amh* and are divided variously among clades of *Sebastes*.

### BLAST queries for ‘usual suspect’ sex determination genes

We wanted to determine the location of ‘usual suspect’ SD genes in our study species and assess whether these genes occurred near any sex-biased regions. To accomplish this, we used command-line BLAST to query FASTA sequences for *ar, dmrt1*, and *sox-3/SRY* genes from three-spined stickleback, and *gsdf* from *Sebastes umbrosus* against all available *Sebastes* genomes.

## Results

### GWAS for sex bias

Three species display significant enrichment of sex bias across large portions of one or more chromosomes: *S. carnatus, S. chrysomelas*, and *S. diaconus* (Fig. 1). These regions contain high enough concentrations of significant markers to be easily detected by our rolling mean of significance. *Sebastes carnatus* and *S. chrysomelas* have a similar concentration of low p-values on chromosome 19, between 0 and ∼8Mbp. The signal across this region was detected by *RADSex* as well as our test of allele frequency and heterozygosity differences (Figure S1-S2).

*Sebastes diaconus* has a very high concentration of sex-biased SNPs on both chromosomes 2 and 12 (Fig. 1). On chromosome 2, significant sex-bias extends from ∼5Mbp to the end of the chromosome at just over 40Mbp and is detectable by RADSex and our allele frequency and heterozygosity metrics (Fig. S3). Similar enrichment of significant sex-bias on chromosome 12 was observed in a region spanning 0 to ∼20Mbp (Fig. S4). This unique pattern male-biased alleles across two chromosomes warranted further investigation.

While most *S. diaconus* males possess the full suite of significantly male-biased alleles on both chromosomes 2 and 12, a single male sample, SRR9968840, lacks these male alleles on the majority of chromosome 2 (Figure 2A). Figure 2B reveals that although males and females differ significantly in allele frequency in these regions, no alleles are unique to one sex.

Alignment to the commonly scaffolded *S. miniatus* genome suggests that sibling species *S. crocotulus* and *S. miniatus* share three narrow regions of sex-bias: ∼33.9Mbp on chromosome 6, ∼8.25Mbp on chromosome 13, and ∼7.30Mbp on chromosome 17 (Fig. 1). A comparison of mean sample depth between sexes in this region revealed significant male bias (Fig. S3).

Only two significant markers in *S. ruberrimus* align to chromosome-level contigs: on chromosomes 4 (36.54 Mbp) and 5 (17.35 Mbp), while four more map to unplaced contigs. These markers are significantly biased toward males; their distribution between sexes, as revealed by RADSex, is visible in supplementary figure 6. Since these markers are not neighboured by other significant markers, as would be expected by a sex determining region, these may represent misplaced reference sequence. No significant sex bias was detected in either *S. pinniger* or *S. paucispinis*.

### Mapping canary, sunset, and vermillion rockfish data to higher quality genomes

While this analysis yielded no novel insight into sex-bias in *S. pinniger*, narrow peaks on an unplaced contig (CAKALS010000047.1) were shared by *S. miniatus* and *S. crocotulus* (Fig. 3A). BLAST results indicate that these are homologous to the peaks previously mapped to RagTag assembly chromosomes 6, 13, and 17. Males had consistently higher read depth along this contig which, considering our likewise analysis of the RagTag alignment, was expected (Fig. 3C; Fig. S5). Another region of sex bias was detected in *S. crocotulus*, along the first megabase of chromosome 24. We found that this differentiation is due to large swathes of missing data in *S. crocotulus* females (Fig. 3B; Fig. S9).

### ssembly of amh phylogeny in Sebastes and their relatives

We found that the *amhy* gene in *S. schlegelii* forms a monophyletic group with copies of *amh* found primarily on chromosome 4, and includes representatives from *Sebastes* sister genus, *Hozukius*. This supports an early origin of *amhy* prior to the split of *Sebastes-Hozukius* but after the divergence of *Sebastiscus* (Fig. 4A).

The initial characterization of *amhy* identifies two insertions in intron 4 and used them as markers to separate the gene copies (Song et al., 2021). By comparing the outgroup sequence in *Sebastolobus*, we found that these are instead deletions in *amh*, rather than insertions in *amhy*. We also found that these deletions are not monomorphic in all orthologs of *amh*, and individual may possess one, both or neither deletion (Fig. 4B).

Our BLAST search results reveal that the locations of orthologous common SD genes are highly conserved within *Sebastes*, and none overlapped regions enriched for sex-biased markers. In our study species, only the *amh* locus in *S. miniatus* lies on a different chromosome from the rest (Fig. 4, Table S1).

## Discussion

By employing an array of analytical approaches, we find evidence of a wide diversity of sex determining mechanisms within just a handful of *Sebastes* species in the Pacific Ocean. Further, the varied metrics permit characterization of the type and degree of differentiation within sex-determining regions. Diversity among SD regions, origins, and mechanisms within a dataset comprising only eight species exemplifies the lability of SD systems in the recently speciated genus (Fig. 5).

**Figure 5.**
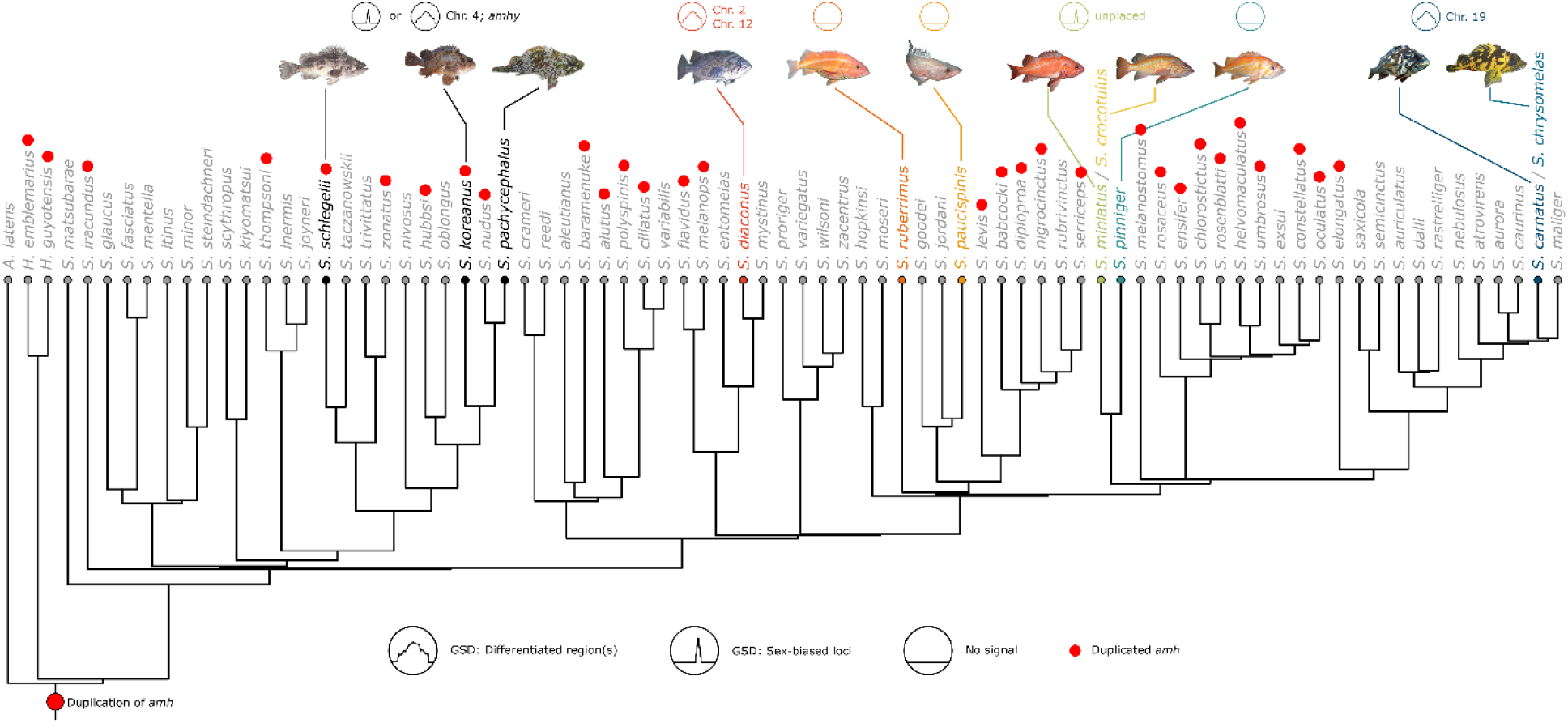
A graphical summary of our findings. Sex-determining regions in *Sebastes* are highly variable in their location and degree of differentiation. The duplication of *amh* predates the *Hozukius*/*Sebastes* split but may not persist in all extant *Sebastes*. Cooption of the duplicated *amhy* as MSD appears to be monophyletic, specific to northwest Pacific species

### Methods to study sex determination

Many different methodological approaches have been employed to identify genomic regions that differ between sexes (Grayson et al., 2022). Depending on the type and degree of differentiation, different analytical methods yield different results. For example, newly differentiating homomorphic sex chromosomes may retain much the same genomic backdrop, their differentiation limited to allele frequency, whereas sex-specific regions are easily recognized by a presence/absence bias or vast differences in sample read depth. We employ three metrics to cover these bases: site allele frequency and heterozygosity inform differentiation before degeneration, whereas presence/absence comparison suggests a deletion in one sex or insertion in the other. Our comprehensive suite of genomic analyses provides both fine-scale resolution of sex-biased markers and in-depth characterization of sex chromosome differentiation.

New methods and programs are constantly in development to address this issue. Here, we compare the effectiveness of one such program (RADSex) to a traditional variant calling pipeline and our various genomic analyses. While RADSex did detect large scale genomic variation in *S. carnatus, S. chrysomelas*, and *S. diaconus*, this high throughput method did not register significant differences at a finer scale in *S. miniatus* and *S. crocotulus*. Our presence/absence metric revealed a much higher density of sex-specific markers at these sites, which, when analyzed for mean sample depth reveal a highly differentiated ∼120kbp contiguous segment of DNA, almost exclusive to males. This difference in resolution is likely due to the way RADSex requires identical reads. In traditional paired-read RADseq, the start of read one is anchored by a restriction digest site but read two are variable, due to random shearing of the other end of the fragment. This means the second reads are unlikely to start at the same position, thus making them non-identical even if they don’t have any genetic polymorphism. As a result, while RADSex only showed two differentiated markers, individually called variants showed that this difference was spread across a larger region, including three peaks in male read depth covered by second reads (Fig. 3C). This gave us more confidence that this represented a true difference rather than an isolated mis-mapping of two reads.

One advantage of RADSex is the identification of sex biased markers is reference-free. Although in our study we have quality reference genomes for all species, that is not always the case. This also provides an advantage in cases where the reference genome is the homogametic sex. In this case, read mapping will be especially poor for reads from the sex determining region (e.g. Y chromosome) since the true mapping location is not present and may lead to spurious sex biased regions across the genome. In our study, the samples used to construct reference genomes were not sexed, therefore we do not know if sex determining regions are missing. From RADSex, we don’t find a large number of unmapped sex biased markers, suggesting that our reference-based analyses are not hampered by this issue.

The advent of streamlined programs to study sex chromosomes provides many useful tools. It is crucial to note, however, that each analytical method comes with a specific set of strengths and weaknesses.

### Two putative sex chromosomes in deacon rockfish

We have identified two putative sex chromosomes in *S. diaconus*: chromosomes 2 and 12. The breadth of these sex-biased regions indicate that the differentiation is in a mature stage. However, many of these markers are still present in some proportion of females (Figure S6). Indeed, the sex-bias on both chromosomes is in allele frequency and heterozygosity, with very few sites specific to one sex (Figures S3-S4).

According to theory on sex chromosome evolution, recombination suppression evolves over time and can result in evolutionary strata along the Y-chromosome (Bergero & Charlesworth, 2009). This gradual accumulation is somewhat in contrast to the large sex chromosome size in *S. diaconus*. Considering this sex determining region is not shared with any of our other tested species, it is likely to have evolved after the origin of *Sebastes* and is perhaps much younger. This raises an interesting question about how a region of this size can evolve in such a short period of time.

Based on the enrichment of male-specific loci (Figure 1), the *S. diaconus* sex chromosome spans most of chromosomes 2 and 12, but there is not a hard barrier on the edges of this region. This means that haplotypes in these neighbouring regions are slightly more likely in one sex or the other, but they are not as linked as the core region. This could because the sex associated region is expanding but has not perfectly suppressed recombination in these outlying regions. Alternatively, the sex associated region may have been introduced as a large, differentiated haplotype and then has undergone some amount of erosion, reducing its size. This scenario has occurred in the ninespine stickleback which has a Y-chromosome derived through introgression with a close relative (Dixon et al., 2019). Most male *S. diaconus* possess at least one copy of the male allele at each of the loci of interest on both sex chromosomes, but one sample lacks these alleles at several sites on chromosome 2. This suggests that the gene(s) most directly responsible for SD in *S. diaconus* reside either on chromosome 12, or in the terminal region of chromosome 2.

It is also interesting that the signal of sex differentiation is shared across two chromosomes. The fact that the sex association signal reaches one end of each chromosome supports that there is a chromosomal translocation that joins these regions. There are examples in nature of large-scale chromosomal rearrangements leading to novel sex chromosomes, including sex chromosome fusions in stickleback (Sardell et al., 2021) and a reciprocal translocation in common frogs, which resulted in two physically separate but coinherited sex chromosomes (Scott, 2019). A third explanation may be found in Chinook salmon, in whom the sex-determining locus is readily translocated between two chromosomes (McKinney et al., 2020). Given the lack of a chromosome scale reference genome for this species, we cannot say if the translocation was involved in the formation of the sex chromosome, or whether an ancestral translocation in this species was then co-opted by a sex determining region, but we consider the first to be more parsimonious.

### Chromosome 19 is the sex chromosome in gopher and black-and-yellow rockfishes

Our analysis identifies chromosome 19 as a putative sex chromosome in the sibling species gopher (*S. carnatus*) and black-and-yellow (*S. chrysomelas*) rockfishes. While the smaller sample sizes (n = 20 in each species) used for study by the original authors somewhat hinder our statistical power, it appears that several markers on this chromosome are male-specific. The differences in presence/absence analyses of both species indicate that *S. carnatus* has a higher degree of heteromorphism (Fig. S1-S2).

These species have a very recent common ancestor (Fowler & Buonaccorsi, 2016) and since they have inherited the same sex chromosome, offer an opportunity to compare differentiation between the species. As in *S. diaconus*, allele frequency and heterozygosity are the main signal of sex differences, indicative of a sex chromosome in the early stages of differentiation. In comparing the two species, we can see slightly more male-specific markers are present in *S. carnatus* than in *S. chrysomelas* (Fig. S1-S2). This may indicate that *S. carnatus* is experiencing a faster rate of sex chromosome differentiation, in which males have acquired insertions or females have undergone deletions of male-associated loci.

The presence of a single sex-biased region restricted to a single chromosome lends support to the classic paradigm of differentiating sex chromosomes in these species. Indeed, the central peak of significant marker density which declines outward in both species is consistent with typical patterns of linkage disequilibrium. Notably, a BLAST query of common SD genes against the *S. carnatus* and *S. chrysomelas* genomes did not provide any hits on chromosome 19, thus we do not know the actual genetic mechanism. This includes *gsdf*, a gene noted by Fowler & Buonaccorsi (2016) as a candidate for cooption as MSD in gopher and black-and-yellow rockfish. Though *gsdf* does reside on chromosome 19 in these species, it is well outside the differentiated region and is therefore not likely to be directly responsible for maleness. The sex of our reference genomes samples is not known; it is possible that the reference genomes for these species are female and so our BLAST query would not find any male-specific genes.

### An unplaced sex-determining region in sunset and vermillion rockfishes

We have identified an unplaced, putative sex-determining region in the sibling species *S. miniatus* and *S. crocotulus*. Differences in mean sample depth reveal distinct peaks of sex differentiation in these species: four in *S. miniatus* and three in *S. crocotulus*. The shape of these peaks however, while suggestive that the degree of male bias tapers outward from a single locus, is an artifact of RADseq technology, where read depth is highest near the first read cut site. All peaks are on a single unplaced contig, and all RADtags on that region have a significant male bias, suggesting the entire contig is a male-specific haplotype.

Mapping of the sunset and vermillion rockfish data to a higher quality reference genome was critical in accurate characterization of the SD region; the RagTag-assembled alignment scattered the signal of sex differentiation onto different chromosomes. Although the reference genomes we used in our initial analyses are at the chromosome level, they were assembled using Illumina short-read data and then scaffolded with a chromosome-scale relative. Under this scenario, it is more likely that individual contigs of a highly diverged SD region are randomly scaffolded due to a lack of obvious synteny.

The significant differentiation observed on chromosome 24 of *S. crocotulus* presents an intriguing possibility: that our unplaced sex-determining region could be at the start of chromosome 24. As we established, both the unplaced SD region and the differentiation along the first megabase of chromosome 24 are characterized by missing data in females. At the start of chromosome 24, there are two distinct haplotypes, one of which is characterized by large amounts of missing data for some SNPs (the “deletion haplotype”). Interestingly, we see some evidence of limited recombination between the haplotypes, which suggests that the haplotypes can recombine, but also that recombination is uncommon (Fig. 3B). We found that while the deletion haplotype is not sex-specific, it does exhibit significant female bias in *S. crocotulus*. Taken together, this supports a scenario where the deletion haplotype is older than the speciation of *S. crocotulus* and *S. miniatus* and has either become recently linked or unlinked to the SDR in one species.

### Weak support for GSD in yelloweye rockfish; no indication of GSD in boccacio and canary rockfish

Significant sex-biased markers are detected in *S. ruberrimus* (Figure S6), but while the markers exhibit significant male bias, none are exclusive to males. Only a single marker could be mapped to chromosome 4, while the others are all on unplaced scaffolds. It is possible that, like *S. miniatus* and *S. crocotulus*, the seemingly disparate sex-biased markers all originate from the same region but were incorrectly scaffolded. Unfortunately, the small number and placement of these markers do not permit further analysis without mapping to a higher quality genome.

No outlying sex-biased markers were detected by our analyses of *S. paucispinis* and *S. pinniger*. As *S. paucispinis* was the species with the fewest samples in this study (n = 13), we have limited confidence in this assessment pertaining to that species.

A lack of genetic markers associated with sex suggests SD by abiotic environmental factors, or social sex ratio pressures. While ESD is perhaps best known in reptiles, it has also been demonstrated in bony fishes, with temperature as the primary environmental determinant of sex (Yamamoto et al., 2019). Temperature has even been shown to affect sex ratios in *S. schlegelii* (Omoto et al., 2010), who (as we mention above) otherwise exhibit demonstrable heterogametic GSD (Song et al., 2021). Other abiotic factors, such as salinity (Saillant et al., 2003) and pH (Reddon Hurd, 2013) have also been shown to induce skewed sex ratios in fishes. This variety of SD systems within a recently speciated clade is not unprecedented; African cichlid fishes exhibit an enormous diversity of SD mechanisms, including several independently evolved sex chromosomes (Gammerdinger Kocher, 2018; El Taher et al., 2021), polygenic GSD (Roberts et al., 2016), ESD (Renn Hurd, 2021), and intraspecific variation in SD mechanisms (Lichilín et al., 2023). Further investigation into the forces underlying ESD in these and other species is warranted, as warming and acidifying oceans leave species whose gonadal differentiation are influenced by environmental factors much more vulnerable to impacts on sex ratios.

### Duplication of amh before speciation of Sebastes

Our analysis found that the duplication of *amh* found by Song et al. (2021) predates the *Hozukius/Sebastes* speciation event. Therefore, we expect this duplication to have the potential to be present in all extant *Sebastes* species. While our phylogeny detects whole gene copies in only 28 *Sebastes* species, in some cases we expect that the duplication is male-specific and would not be detectable on genomes assembled from female samples. As Kolora et al. (2021) did not sex their samples before assembly, we are unable to make that determination. Further, when the duplicated *amhy* in *S. schlegelii* may have acquired its SD function remains in question. Song et al. (2021) showed that males and females differed in *amh* copy number in *S. schlegelii, S. koreanus* and the closely related *S. pachycephalus*, suggesting perhaps that this could be shared amongst all northwest Pacific rockfish.

There are two duplications in *Sebastes* that stand out as evidence for the lability of SD genes in this genus. In *Sebastes reedi*, the entire sequence of *amh* on chromosome 5 is found in an inverted duplication immediately adjacent to the original gene copy. *Sebastes ensifer* possesses two copies of *amhy*, one in the typical chromosome 4 position and one on chromosome 5, the typical position for *amh*. This may suggest a translocation or gene conversion that overwrote the original *amh*, although we cannot rule out that is a bioinformatic artifact during assembly.

### Areas for further research

We have demonstrated that SD among northeast Pacific *Sebastes* is highly diverse and, due to the relatively undifferentiated sex chromosomes we uncovered, highly labile. However, as our study comprises only eight species, our evolutionary inferences are limited. Further studies involving sexed sequence data from geographically diverse *Sebastes* may shed light on the evolutionary trajectory of SD systems in this genus. This could reveal shared mechanisms of GSD, gain-of-function events, translocations, and other genomic rearrangements underpinning the lability of SD.

Deeper investigation into the genes underlying SD is required. While our study has discovered sex chromosomes in three species, and potential SD loci in two more, the precise genetic mechanisms at work are yet unknown. Until recently, most eukaryotic genomes sequences published were haploid representations of a diploid individual. For species with partially differentiated SD regions, the SD region is often shunted to an unplaced scaffold. Fully phased genomes for the heterogametic sex allows for appropriate read mapping and an accurate assessment of gene level differences in the SD region (Carey et al., 2022). Future studies providing complete chromosome-level reference assemblies of different rockfish species will greatly aid in the identification of SD loci in the *Sebastes* clade.

## Conclusion

Sex determination systems in nature are myriad and dynamic. While patterns do emerge in the evolution of SD mechanisms in animals, ever more analyses of systems without fixed modes of SD challenge the classic paradigm underlying sex chromosome differentiation. Here, we demonstrate that *Sebastes* rockfish exhibit a wide diversity of SD mechanisms of independent origins. In this recently speciated genus, we find evidence for heterogametic, multi-locus, and environmental SD mechanisms, in only a handful of study species.

Further study into the diversity of SD in this highly speciose genus is essential to enhance our understanding of the selective and stochastic forces underlying sex determination. Indeed, our results contribute to a growing body of evidence that our current understanding of SD evolution is lacking. As studies into previously overlooked taxa reveal the wealth and lability of SD systems in nature, we stand to better characterize the genetic and ecological drivers of sex differentiation, a fundamental trait spanning the tree of life.

## Acknowledgements

We’d like to thank all the rockfish researchers whose data was used in this project: Kelly S. Andrews, Benjamin L.S. Fowler, Gary C. Longo, Krista M. Nichols, Kathleen G. O’Malley, Daniel M. Tonnes, Felix Vaux, and the late Vincent P. Buonaccorsi. Given the scarcity of sexed sequence data, we are very grateful for your efforts. We are also grateful to the underwater photographers whose original work was cropped for our figure, including Rick Ramsey and K. Lynne Yamanaka. This work was supported by an NSERC discovery grant to Gregory L. Owens.

## References

Alonge, M., Lebeigle, L., Kirsche, M., Aganezov, S., Wang, X., Lippman, Z. B., … Soyk, S. (2021). Automated assembly scaffolding elevates a new tomato system for high-throughput genome editing. bioRxiv. doi: 10.1101/2021.11.18.469135

Andrews, K. S., Nichols, K. M., Elz, A., Tolimieri, N., Harvey, C. J., Pacunski, R., … Tonnes, D. M. (2018). Cooperative research sheds light on population structure and listing status of threatened and endangered rockfish species. Conservation Genetics, 19(4), 865–878. doi: 10.1007/s10592-018-1060-0

Bachtrog, D. (2013). Y-chromosome evolution: Emerging insights into processes of Y-chromosome degeneration. Nature Reviews Genetics, 14(2), 113–124. doi: 10.1038/nrg3366

Bachtrog, D., Mank, J. E., Peichel, C. L., Kirkpatrick, M., Otto, S. P., Ashman, T.-L., … The Tree of Sex Consortium (2014). Sex Determination: Why So Many Ways of Doing It? PLOS Biology, 12(7), e1001899. doi: 10.1371/journal.pbio.1001899

Bergero, R., Charlesworth, D. (2009). The evolution of restricted recombination in sex chromosomes. Trends in Ecology Evolution, 24(2), 94–102. https://doi.org/10.1016/j.tree.2008.09.010

Broad Institute. (2019). Picard Toolkit. Broad Institute, GitHub Repository. Retrieved from https://github.com/broadinstitute/picard

Camacho, C., Coulouris, G., Avagyan, V., Ma, N., Papadopoulos, J., Bealer, K., Madden, T. L. (2009). BLAST+: Architecture and applications. BMC Bioinformatics, 10(1), 421. doi: 10.1186/1471-2105-10-421

Carey, S. B., Lovell, J. T., Jenkins, J., Leebens-Mack, J., Schmutz, J., Wilson, M. A., Harkess, A. (2022). Representing sex chromosomes in genome assemblies. Cell Genomics, 2(5), 100132. doi: 10.1016/j.xgen.2022.100132

Charlesworth, B., Harvey, P. H., Charlesworth, D. (2000). The degeneration of Y chromosomes. Philosophical Transactions of the Royal Society of London. Series B: Biological Sciences, 355(1403), 1563–1572. doi: 10.1098/rstb.2000.0717

Chiba, S. N., Ohashi, S., Tanaka, F., Suda, A., Fujiwara, A., Snodgrass, D., … Suzuki, N. (2021). Effectiveness and potential application of sex-identification DNA markers in tunas. Marine Ecology Progress Series, 659, 175–184. doi: 10.3354/meps13563

Couger, M. B., Roy, S. W., Anderson, N., Gozashti, L., Pirro, S., Millward, L. S., Kim, M., Kilburn, D., Liu, K. J., Wilson, T. M., Epps, C. W., Dizney, L., Ruedas, L. A., Campbell, P. (2021). Sex chromosome transformation and the origin of a male-specific X chromosome in the creeping vole. Science, 372(6542), 592–600. doi: 10.1126/science.abg7019

Danecek, P., Bonfield, J. K., Liddle, J., Marshall, J., Ohan, V., Pollard, M. O., … Li, H. (2021). Twelve years of SAMtools and BCFtools. GigaScience, 10(2), giab008. doi: 10.1093/gigascience/giab008

Dixon, G., Kitano, J., Kirkpatrick, M. (2019). The Origin of a New Sex Chromosome by Introgression between Two Stickleback Fishes. Molecular Biology and Evolution, 36(1), 28–38. https://doi.org/10.1093/molbev/msy181

Edgar, R. C. (2004). MUSCLE: Multiple sequence alignment with high accuracy and high throughput. Nucleic Acids Research, 32(5), 1792–1797. doi: 10.1093/nar/gkh340

Edgecombe, J., Urban, L., Todd, E. V., Gemmell, N. J. (2021). Might Gene Duplication and Neofunctionalization Contribute to the Sexual Lability Observed in Fish? Sexual Development, 15(1–3), 122–133. doi: 10.1159/000515425

Edvardsen, R. B., Wallerman, O., Furmanek, T., Kleppe, L., Jern, P., Wallberg, A., … Rubin, C.-J. (2022). Heterochiasmy and the establishment of gsdf as a novel sex determining gene in Atlantic halibut. PLOS Genetics, 18(2), e1010011. doi: 10.1371/journal.pgen.1010011

El Taher, A., Ronco, F., Matschiner, M., Salzburger, W., Böhne, A. (2021). Dynamics of sex chromosome evolution in a rapid radiation of cichlid fishes. Science Advances, 7(36), eabe8215. doi: 10.1126/sciadv.abe8215

Feron, R., Pan, Q., Wen, M., Imarazene, B., Jouanno, E., Anderson, J., … Guiguen, Y. (2021). RADSex: A computational workflow to study sex determination using restriction site-associated DNA sequencing data. Molecular Ecology Resources, 21(5), 1715–1731. doi: 10.1111/1755-0998.13360

Fowler, B. L. S., Buonaccorsi, V. P. (2016). Genomic characterization of sex-identification markers in Sebastes carnatus and Sebastes chrysomelas rockfishes. Molecular Ecology, 25(10), 2165–2175. doi: 10.1111/mec.13594

Gammerdinger, W. J., Kocher, T. D. (2018). Unusual Diversity of Sex Chromosomes in African Cichlid Fishes. Genes, 9(10), Article 10. https://doi.org/10.3390/genes9100480

Garrison, E., Marth, G. (2012). Haplotype-based variant detection from short-read sequencing. arXiv. doi: 10.48550/arXiv.1207.3907

Gessler, D. D. G. (1995). The constraints of finite size in asexual populations and the rate of the ratchet. Genetics Research, 66(3), 241–253. doi: 10.1017/S0016672300034686

Grayson, P., Wright, A., Garroway, C. J., Docker, M. F. (2022, February 22). SexFindR: A computational workflow to identify young and old sex chromosomes. bioRxiv. doi: 10.1101/2022.02.21.481346

Hall, T. A. (1999). BioEdit: A user-friendly biological sequence alignment editor and analysis program for Windows 95/98/NT. Nucleic Acids Symposium Series, 41, 95–98.

Hattori, R. S., Strüssmann, C. A., Fernandino, J. I., Somoza, G. M. (2013). Genotypic sex determination in teleosts: Insights from the testis-determining amhy gene. General and Comparative Endocrinology, 192, 55–59. doi: 10.1016/j.ygcen.2013.03.019

Jeffries, D. L., Mee, J. A., Peichel, C. L. (2022). Identification of a candidate sex determination gene in Culaea inconstans suggests convergent recruitment of an Amh duplicate in two lineages of stickleback. Journal of Evolutionary Biology, n/a(n/a). doi: 10.1111/jeb.14034

Kolora, S. R. R., Owens, G. L., Vazquez, J. M., Stubbs, A., Chatla, K., Jainese, C., … Sudmant, P. H. (2021). Origins and evolution of extreme life span in Pacific Ocean rockfishes. Science, 374(6569), 842–847. doi: 10.1126/science.abg5332

Kong, A., Thorleifsson, G., Gudbjartsson, D. F., Masson, G., Sigurdsson, A., Jonasdottir, A., … Stefansson, K. (2010). Fine-scale recombination rate differences between sexes, populations and individuals. Nature, 467(7319), 1099–1103. doi: 10.1038/nature09525

Kratochvíl, L., Stöck, M., Rovatsos, M., Bullejos, M., Herpin, A., Jeffries, D. L., … Pokorná, M. J. (2021). Expanding the classical paradigm: What we have learnt from vertebrates about sex chromosome evolution. Philosophical Transactions of the Royal Society B: Biological Sciences, 376(1833), 20200097. doi: 10.1098/rstb.2020.0097

Li, X.-Y., Gui, J.-F. (2018). Diverse and variable sex determination mechanisms in vertebrates. Science China Life Sciences, 61(12), 1503–1514. doi: 10.1007/s11427-018-9415-7

Lichilín, N., Salzburger, W., Böhne, A. (2023). No evidence for sex chromosomes in natural populations of the cichlid fish Astatotilapia burtoni. G3 Genes|Genomes|Genetics, jkad011. https://doi.org/10.1093/g3journal/jkad011

Liu, X., Dai, S., Wu, J., Wei, X., Zhou, X., Chen, M., … Wang, D. (2022). Roles of anti-Müllerian hormone and its duplicates in sex determination and germ cell proliferation of Nile tilapia. Genetics, 220(3), iyab237. doi: 10.1093/genetics/iyab237

Longo, G. C., Harms, J., Hyde, J. R., Craig, M. T., Ramón-Laca, A., Nichols, K. M. (2022). Genome-wide markers reveal differentiation between and within the cryptic sister species, sunset and vermilion rockfish. Conservation Genetics, 23(1), 75–89. doi: 10.1007/s10592-021-01397-4

Luckenbach, J. A., Borski, R. J., Daniels, H. V., Godwin, J. (2009). Sex determination in flatfishes: Mechanisms and environmental influences. Seminars in Cell Developmental Biology, 20(3), 256–263. doi: 10.1016/j.semcdb.2008.12.002

Mangel, M., Kindsvater, H. K., Bonsall, M. B. (2007). Evolutionary Analysis of Life Span, Competition, and Adaptive Radiation, Motivated by the Pacific Rockfishes (Sebastes). Evolution, 61(5), 1208–1224. doi: 10.1111/j.1558-5646.2007.00094.x

Mank, J. E., Promislow, D. E. L., Avise, J. C. (2006). Evolution of alternative sex-determining mechanisms in teleost fishes. Biological Journal of the Linnean Society, 87(1), 83–93. doi: 10.1111/j.1095-8312.2006.00558.x

Marshall Graves, J. A., Peichel, C. L. (2010). Are homologies in vertebrate sex determination due to shared ancestry or to limited options? Genome Biology, 11(4), 205. doi: 10.1186/gb-2010-11-4-205

McKinney, G. J., Nichols, K. M., Ford, M. J. (2021). A mobile sex-determining region, male-specific haplotypes and rearing environment influence age at maturity in Chinook salmon. Molecular Ecology, 30(1), 131–147. doi: 10.1111/mec.15712

Muller, H. J. (1932). Some Genetic Aspects of Sex. The American Naturalist, 66(703), 118–138. doi: 10.1086/280418

Muller, H. J. (1964). The relation of recombination to mutational advance. Mutation Research/Fundamental and Molecular Mechanisms of Mutagenesis, 1(1), 2–9. doi: 10.1016/0027-5107(64)90047-8

Nakamoto, M., Uchino, T., Koshimizu, E., Kuchiishi, Y., Sekiguchi, R., Wang, L., … Sakamoto, T. (2021). A Y-linked anti-Müllerian hormone type-II receptor is the sex-determining gene in ayu, Plecoglossus altivelis. PLOS Genetics, 17(8), e1009705. doi: 10.1371/journal.pgen.1009705

Nguyen, L.-T., Schmidt, H. A., von Haeseler, A., Minh, B. Q. (2015). IQ-TREE: A Fast and Effective Stochastic Algorithm for Estimating Maximum-Likelihood Phylogenies. Molecular Biology and Evolution, 32(1), 268–274. doi: 10.1093/molbev/msu300

Ohno, S. (1967). Sex Chromosomes and Sex-linked Genes. New York: Springer-Verlag.

Pokorná, M., Kratochvíl, L. (2009). Phylogeny of sex-determining mechanisms in squamate reptiles: Are sex chromosomes an evolutionary trap? Zoological Journal of the Linnean Society, 156(1), 168–183. doi: 10.1111/j.1096-3642.2008.00481.x

Rambaut, A. (2018). FigTree. Retrieved from tree.bio.ed.ac.uk/software/figtree/

Reddon, A. R., Hurd, P. L. (2013). Water pH during early development influences sex ratio and male morph in a West African cichlid fish, Pelvicachromis pulcher. Zoology, 116(3), 139–143. doi: 10.1016/j.zool.2012.11.001

Renn, S. C. P., Hurd, P. L. (2021). Epigenetic Regulation and Environmental Sex Determination in Cichlid Fishes. Sexual Development, 15(1–3), 93–107. https://doi.org/10.1159/000517197

Renner, S. S., Ricklefs, R. E. (1995). Dioecy and its correlates in the flowering plants. American Journal of Botany, 82(5), 596–606. doi: 10.1002/j.1537-2197.1995.tb11504.x

Roberts, N. B., Juntti, S. A., Coyle, K. P., Dumont, B. L., Stanley, M. K., Ryan, A. Q., Fernald, R. D., Roberts, R. B. (2016). Polygenic sex determination in the cichlid fish Astatotilapia burtoni. BMC Genomics, 17(1), 835. https://doi.org/10.1186/s12864-016-3177-1

Saillant, E., Fostier, A., Haffray, P., Menu, B., Chatain, B. (2003). Saline preferendum for the European sea bass, Dicentrarchus labrax, larvae and juveniles: Effect of salinity on early development and sex determination. Journal of Experimental Marine Biology and Ecology, 287(1), 103–117. doi: 10.1016/S0022-0981(02)00502-6

Sardell, J. M., Josephson, M. P., Dalziel, A. C., Peichel, C. L., Kirkpatrick, M. (2021). Heterogeneous Histories of Recombination Suppression on Stickleback Sex Chromosomes. Molecular Biology and Evolution, 38(10), 4403–4418. doi: 10.1093/molbev/msab179

Scott, M. F. (2019). Causes and consequences of reciprocal translocations on sex chromosomes. Molecular Ecology, 28(8), 1863–1865. doi: 10.1111/mec.15064

Song, W., Xie, Y., Sun, M., Li, X., Fitzpatrick, C. K., Vaux, F., … He, Y. (2021). A duplicated amh is the master sex-determining gene for Sebastes rockfish in the Northwest Pacific. Open Biology, 11(7), 210063. doi: 10.1098/rsob.210063

Sunobe, T., Sado, T., Hagiwara, K., Manabe, H., Suzuki, T., Kobayashi, Y., … Miya, M. (2017). Evolution of bidirectional sex change and gonochorism in fishes of the gobiid genera Trimma, Priolepis, and Trimmatom. The Science of Nature, 104(3), 15. doi: 10.1007/s00114-017-1434-z

Tange, O. (2011, February). GNU Parallel—The command-line power tool. The USENIX Magazine, 42–47.

Vasimuddin, Md., Misra, S., Li, H., Aluru, S. (2019). Efficient Architecture-Aware Acceleration of BWA-MEM for Multicore Systems. 2019 IEEE International Parallel and Distributed Processing Symposium (IPDPS), 314–324. doi: 10.1109/IPDPS.2019.00041

Vaux, F., Rasmuson, L. K., Kautzi, L. A., Rankin, P. S., Blume, M. T. O., Lawrence, K. A., … O‘Malley, K. G. (2019). Sex matters: Otolith shape and genomic variation in deacon rockfish (Sebastes diaconus). Ecology and Evolution, 9(23), 13153–13173. doi: 10.1002/ece3.5763

Wright, A. E., Dean, R., Zimmer, F., Mank, J. E. (2016). How to make a sex chromosome. Nature Communications, 7(1), 12087. doi: 10.1038/ncomms12087

Yamamoto, Y., Hattori, R. S., Patiño, R., Strüssmann, C. A. (2019). Chapter Two - Environmental regulation of sex determination in fishes: Insights from Atheriniformes. In B. Capel (Ed.), Current Topics in Developmental Biology (pp. 49–69). Academic Press. doi: 10.1016/bs.ctdb.2019.02.003

