## Supplementary Figures for "Rapid turnover and evolution of sex-determining regions among *Sebastes* rockfishes" (from another version of this preprint: version 1, 2023-01-20)

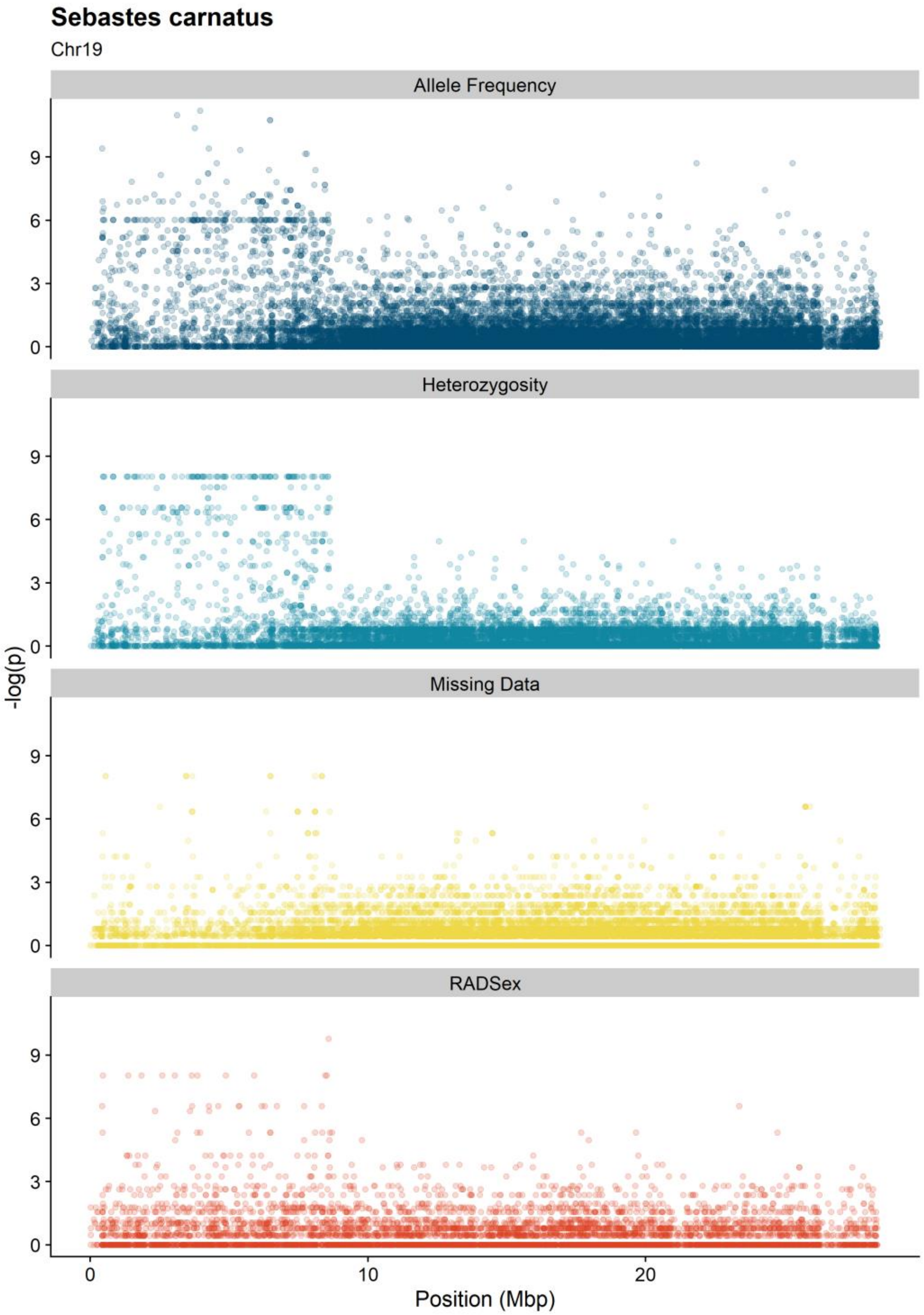


**Figure S1** | Sebastes carnatus putative sex chromosome 19 shows high degrees of differentiation in allele frequency and heterozygosity. Fewer markers in the missing data metric suggest few alleles have yet been purged from the female genotype.

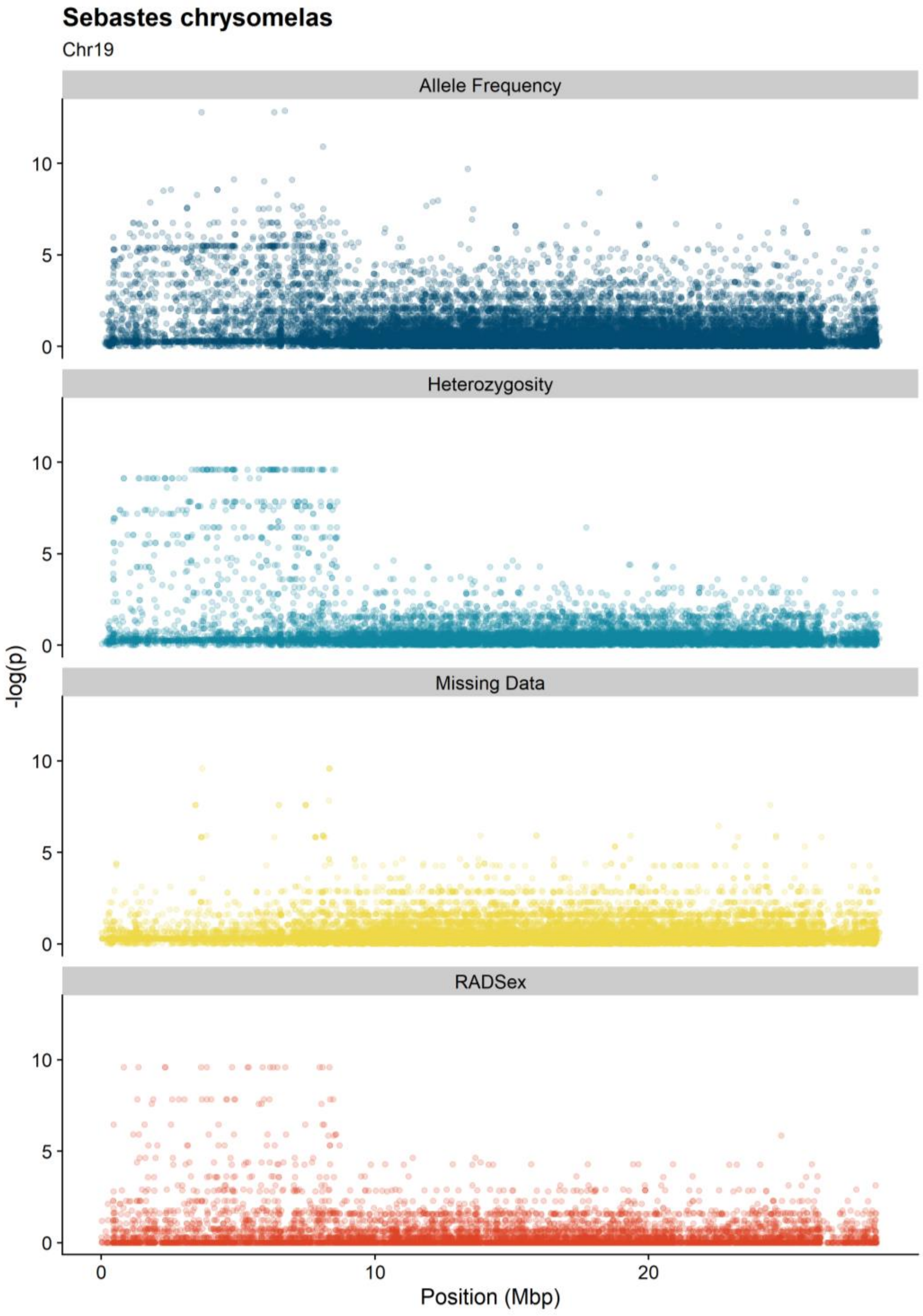


**Figure S2** | Sebastes chrysomelas putative sex chromosome 19 shows high degrees of differentiation in allele frequency and heterozygosity. Fewer markers in the missing data metric suggest few alleles have yet been purged from the female genotype.

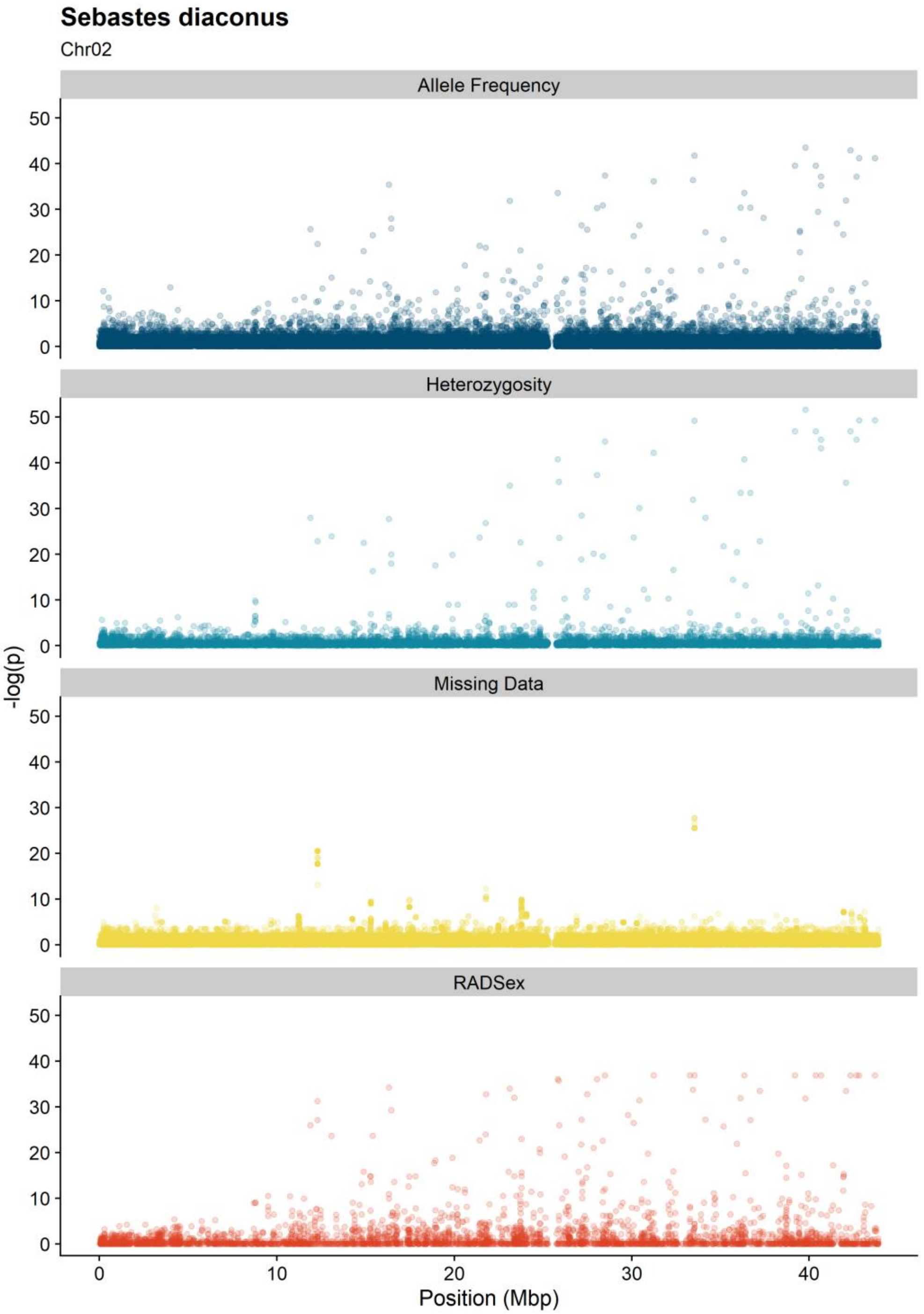


**Figure S3** | Sebastes diaconus putative sex chromosome 02 shows high degrees of differentiation in allele frequency and heterozygosity. Very few markers in the missing data metric suggest few alleles antagonistic to female development.

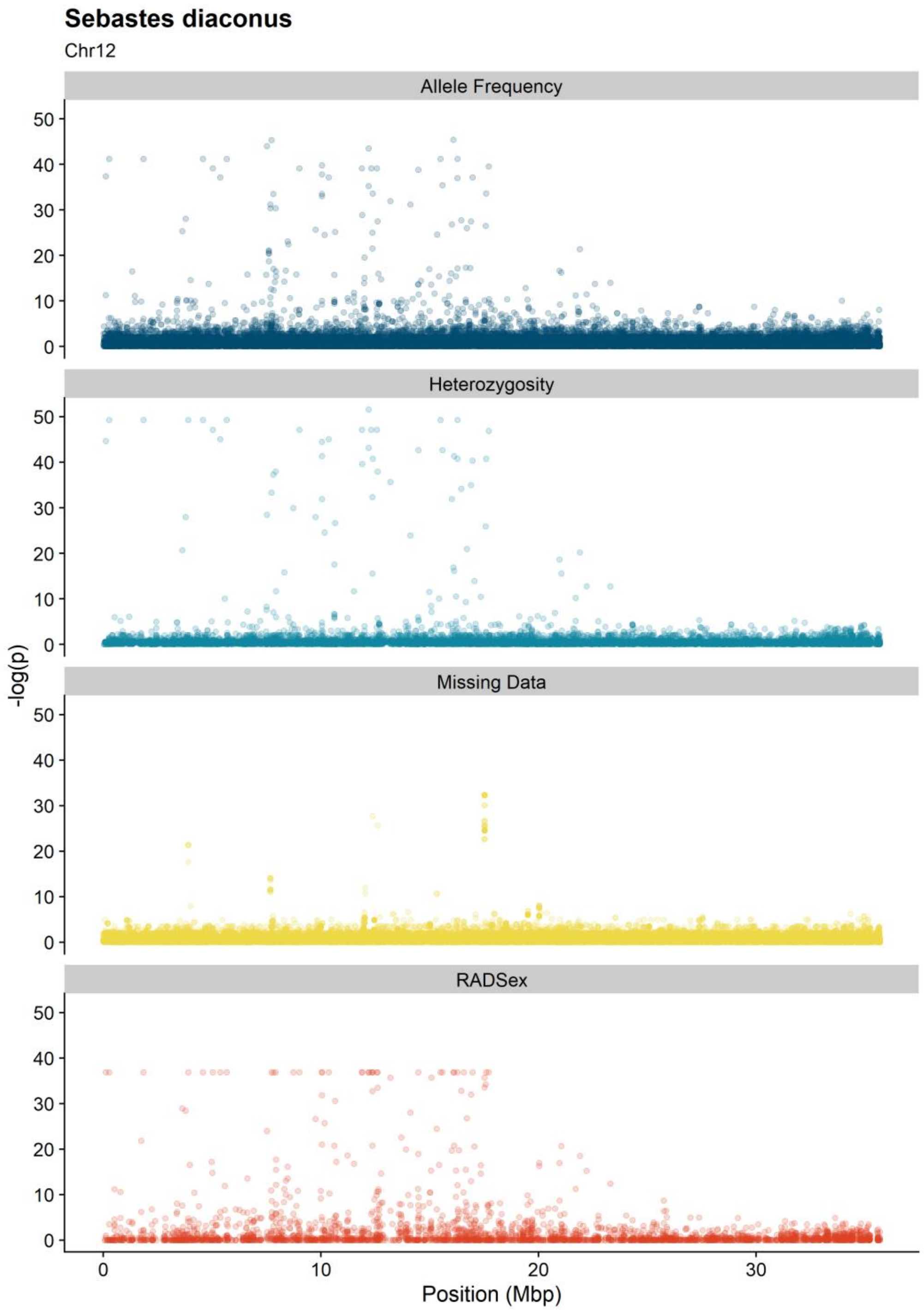


**Figure S4** | Sebastes diaconus putative sex chromosome 12 shows high degrees of differentiation in allele frequency and heterozygosity. Markers in highly specific regions of the missing data metric suggest that some degeneration is underway.

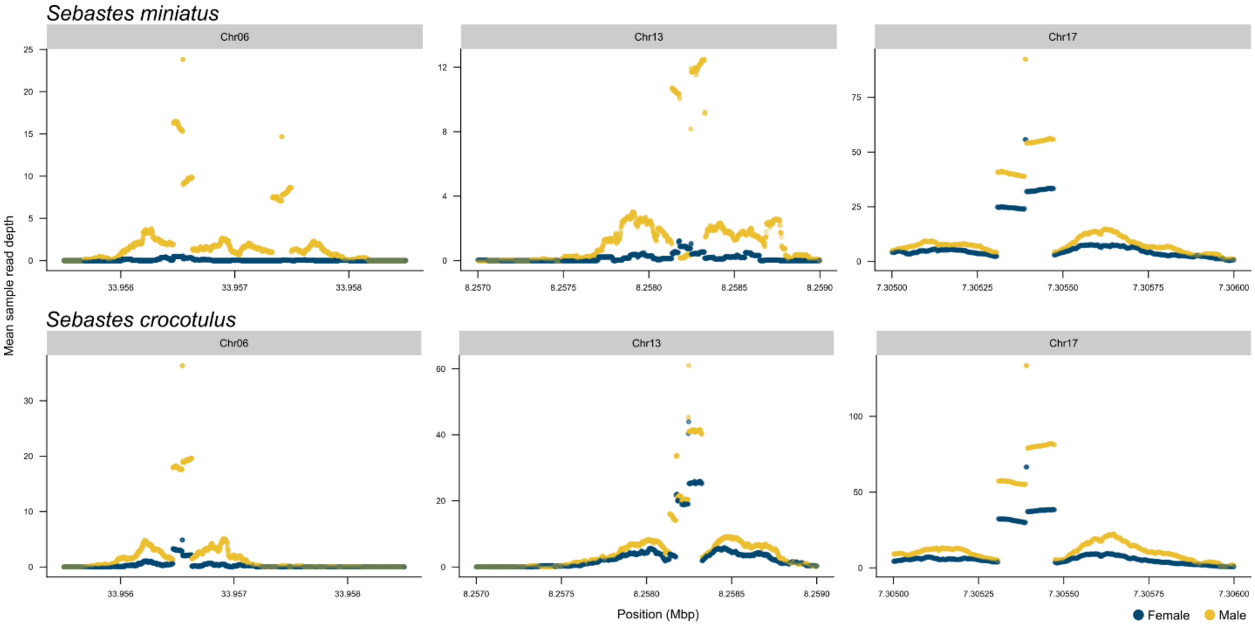


**Figure S5** | Comparison of mean read depth between sexes at three loci in *S. miniatus* and *S. crocotulus.* Common scaffolding to the *S. aleutianus* reference genome scattered the signal of sex bias across three chromosomes. Alignment to a higher quality genome revealed that all three loci map to a single unplaced contig.

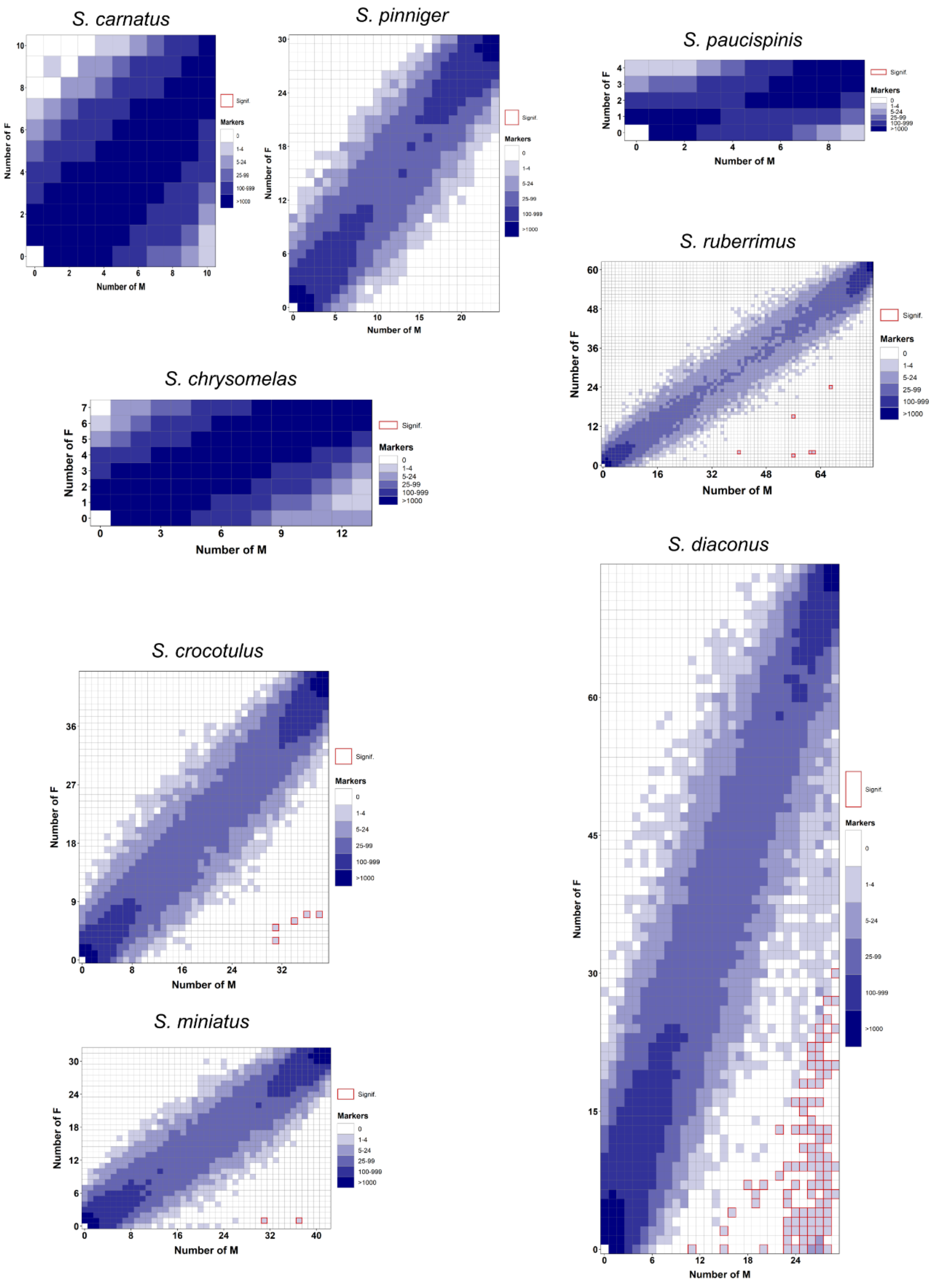


**Figure S6** | Distribution of markers between sexes, computed and visualized by RADSex/sgtr. No markers show significant bias toward females.

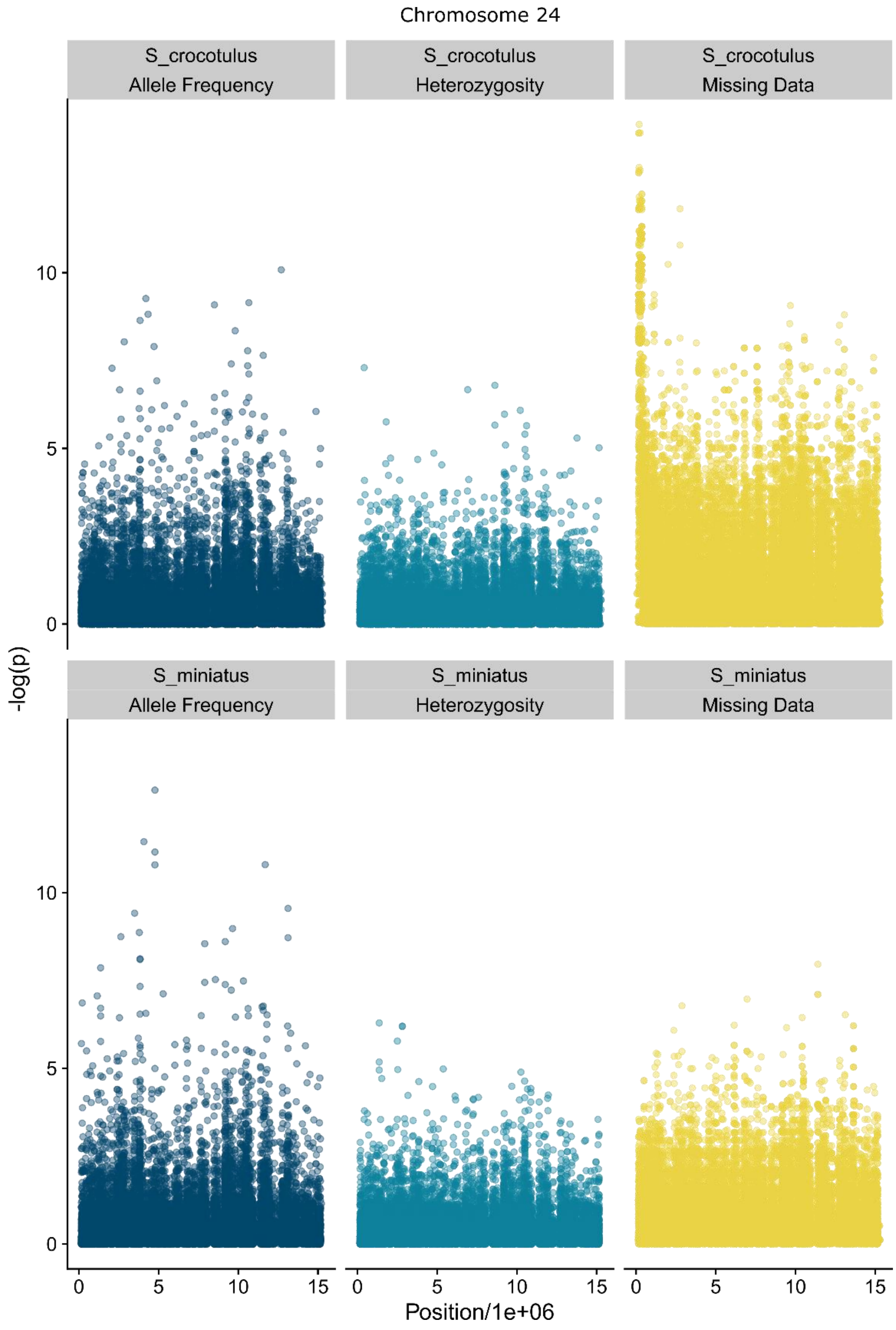


**Figure S7** | Alignment of *S. miniatus* and *S. crocotulus* data to a high-quality genome reveal a sex bias along the first megabase of chromosome 24 in *S. crocotulus*; females have significant amounts of missing data along this region.

| Species | amh | α-ar | α(1A)-ar | α(2A)-ar | α(2C)-ar | α(2DB)-ar | β-ar | β(1)-ar | β(3A)-ar | dmrt1 | sox3/SRY |
|---|---|---|---|---|---|---|---|---|---|---|---|
| *S. carnatus* | Chr05 | Chr10 | Chr10 | Chr03 | Chr03 | Chr10 | Chr17 | Chr21 | Chr06 | *Chr02, *Chr06 | Chr10 |
| *S. diaconus* | Chr05 | Chr10 | Chr10 | Chr03 | Chr03 | Chr10 | Chr17 | Chr21 | Chr06 | *Chr09, *Chr13 | Chr10 |
| *S. miniatus* | Chr20 | Chr10 | Chr10 | Chr03 | Chr03 | Chr10 | Chr17 | Chr21 | Chr06 | *Chr06, *Chr16 | Chr10 |
| *S. paucispinis* | Chr05 | Chr10 | Chr10 | Chr03 | Chr03 | Chr10 | Chr17 | Chr21 | Chr06 | *Chr01, *Chr06 | Chr10 |
| *S. pinniger* | Chr05 | Chr10 | Chr10 | Chr03 | Chr03 | Chr10 | Chr17 | Chr21 | Chr06 | *Chr06, *Chr16 | Chr10 |
| *S. ruberrimus* | Chr05 | Chr10 | Chr10 | Chr03 | Chr03 | Chr10 | Chr17 | Chr21 | Chr06 | *Chr11, *Chr16 | Chr10 |

**Table S1** | Summary of 'usual suspect' gene locations on genome of study species. Only species with a reference genome shown (excludes *S. crocotulus* and *S. chrysomelas*). Asterisks indicate that only short, highly repetitive gene segments registered as hits during the query.
